# Transient receptor potential melastatin 4 (TRPM4) regulates hilar mossy cell loss in temporal lobe epilepsy

**DOI:** 10.1101/2022.10.31.514477

**Authors:** Laura Mundrucz, Angéla Kecskés, Nóra Henn-Mike, Péter Kóbor, Péter Buzás, Rudi Vennekens, Miklós Kecskés

## Abstract

Mossy cells comprise a large fraction of excitatory neurons in the hippocampal dentate gyrus and their loss is one of the major hallmarks of temporal lobe epilepsy (TLE). The vulnerability of mossy cells in TLE is well known in animal models as well as in patients, however the mechanisms leading to cellular death is unclear. One possible explanation for their sensitivity is linked to their specific ion channel composition. TRPM4 is a Ca^2+^-activated non-selective cation channel regulating diverse physiological function of excitable cells. Here, we identified that TRPM4 is present and functionally active in hilar mossy cells. Furthermore, we showed that TRPM4 contributes to mossy cells death following status epilepticus and therefore modulates seizure susceptibility and epilepsy-related memory deficits in the chronic phase of TLE.

## Introduction

Temporal lobe epilepsy (TLE) is a life-threating neurological disorder characterized by recurrent seizures and an increased risk for a wide range of cognitive diseases[1]. Anti-epileptic drugs are ineffective in about 30% of patients demonstrating the urgent need to better understand the pathological mechanisms underlying epileptic seizures[2].

Partial loss of the hilar mossy cells (MC) of the hippocampus is one of the main hallmarks of TLE[3]. MCs are key excitatory neurons of the hippocampus innervating the granule cells (GC) of the dentate gyrus as well as GABAergic interneurons of the hilus[4]. They receive their major excitatory input from GCs via giant synaptic buttons onto large spine complexes along their proximal dendrites[5]. This results in a complex feedback microcircuit: MCs can directly excite GCs as well as indirectly inhibit them via GABAergic interneurons[6]. The opposing effects of MC activation led to debates whether the net effect of MC activation to GCs is inhibitory or excitatory in physiological conditions as well as in epilepsy[7]. Recently, it has been suggested that MC activation can result in a net inhibitory effect on the hippocampus indicating that MC loss in TLE can be an important mechanism leading to recurrent seizures[8]. However, another recent study found that selective MC inhibition decreases status epilepticus severity[9]. Despite the often controversial results about the exact role of MCs in hippocampal excitability their extreme vulnerability in TLE has been well-established in rodent models as well as in human patients[10].

Indeed, one of the most exciting questions about MCs is the reason for their vulnerability in TLE. A possible mechanism suggests that an initial insult resulting in over-excitation of the local circuits presynaptic to MCs increases intracellular calcium and sodium up to a toxic level[11]. Another potential explanation is related to the intrinsic electrophysiological properties of the MCs: these cells are characterized by a low threshold for action potential, weak repolarization with practically no after hyperpolarization (AHP) and broad action potential (AP)[12]. Therefore, a strong and long-lasting excitatory input can easily lead to over excitation, depolarization block and cellular death[3]. Clearly, the unique ion channel expression profile of the MCs leading to the above detailed electrophysiological properties might explain both hypotheses, yet the exact mechanism remains uncertain.

It has been shown already that calcium activated depolarizing currents are responsible for action potential duration and repolarization dynamics in other excitable cell types[13][14]. The ion channels carrying this particular current belongs to the superfamily of Transient Receptor Potential (TRP) ion channels, called Transient Receptor Potential Melastatin 4 (TRPM4) and Transient Receptor Potential Melastatin 5 (TRPM5)[15].

In neurons TRPM4-dependent membrane depolarization can support bursts of action potentials as it was shown in pre-Botzinger complex neurons[16][17]. Furthermore, it can also mediate axonal and neuronal degeneration and cellular death in the animal model of experimental autoimmune encephalomyelitis and multiple sclerosis[18]. In line with the above-mentioned properties, we hypothesized that TRPM4 might play a role in the physiological function of MCs, as well as in the neuronal loss seen in TLE. However, until now there was no information available about the expression and function of TRPM4 on MCs of the dentate gyrus.

Here, with a battery of histological, electrophysiological and behavioral experiments we demonstrate for the first time that (i.) *Trpm4* is present in hilar MC, (ii.) plays a role in their intrinsic electrophysiological properties, (iii.) contributes to MC death following status epilepticus (SE) and therefore (iv.) modulates seizure susceptibility and (v.) epilepsy-related memory deficits in the chronic phase of experimental TLE. Altogether our results shed light on a previously unrecognized role of TRPM4 in neuronal excitability both in healthy and overexcited hippocampus.

## Materials and methods

### Experimental animals

Experiments were performed on 3- to 6-month-old male C57BL/6 N mice and age matched *Trpm4*^*−/−*^ mice. Mice were housed with a 12-h light/12-h dark cycle and allowed water and standard food ad libitum. All animal experiments were performed according to the European Community Council Directive and approved by the local ethics committee (Ethics Committee on Animal Research of Pécs, Hungary, BA02/2000-10/2020 and BA02/2000-56/2022).

### Epilepsy induction and monitoring

Kainic acid (100 nL, 5 mM in saline) was stereotaxically injected into the left hippocampus (2.0 mm posterior, 1.05 mm left, and 1.6 mm ventral to Bregma) using 1μL pipette (Hamilton, 25g needle) under isoflurane anesthesia to induce status epilepticus (SE). SE induction was considered successful if grade 3 or higher seizures were observed based on the Racine’s scale. Mice showing no sign of SE were excluded from the subsequent experiments. Control mice were injected with saline in the same position. 7-10 days after KA or saline injection animals were stereotaxically implanted with two recording electrodes (tungsten, 0.05 mm, insulated, GoodFellow) (2.0 mm posterior, 1.05 mm left/right, and 1.6 mm ventral to Bregma) into both hippocampi, a reference (somatosensory cortex, 2.0 mm posterior, 2.3 mm left, and 0.5 mm ventral to bregma) and a ground electrode (under the skin). Tungsten wires were soldered to a custom made 6 pin connector suitable to connect with the preamplifier. This connector was fixed into the scalp of the mouse using dental cement. After recovery from the implant procedure, mice underwent videoEEG monitoring. During recordings each mouse was individually placed into a 25 cm × 25 cm transparent cage. Same bedding, food and water was supplied as during standard housing. For seizure detection a preamplifier was inserted into the 6-pin connector via a multichannel commutator (Moflon Technology LTD, MC190). This system allowed the mice to freely move throughout the entire cage. EEG signal was acquired at 1 kHZ and band pass filtered at 1.6-2000 Hz (Supertech BioAmp, AD Instruments PowerLab, MultiChannel Systems, W2100). Data were acquired and analyzed using LabChart software (AD Instruments) and Brainstorm[19]. Synchronous video recordings were captured using a webcam (Alcor AWC1080)

### Slice preparation

*In vitro* patch clamp recordings were performed in acute horizontal brain slices taken from C57BL/6N and *Trpm4*^*−/−*^ mice. Under deep isoflurane anesthesia, mice were decapitated and coronal slices (300 μm thick) were cut using a vibratome (Leica VS1200) in ice-cold external solution containing (in mM): 93 NMDG, 2.5 KCl, 25 Glucose, 20 Hepes, 1.2 NaH2PO_4_, 10 MgSO_4_, 0.5 CaCl_2_, 30 NaHCO_3_, 5 L-ascorbate, 3 Na-Pyruvate, 2 thiourea, bubbled with 95% O_2_ and 5% CO_2_. Slices were transferred to artificial cerebrospinal fluid (ACSF) containing (in mM) 2.5 KCl, 10 glucose, 126 NaCl, 1.25 NaH_2_PO_4_, 2 MgCl_2_, 2 CaCl_2_, 26 NaHCO_3_ bubbled with 95% O_2_ and 5% CO_2_. After an incubation period of 10 min at 34 °C in the first solution, the slices were maintained at 20–22 °C in ACSF until use. After recordings, the sections were immersed into fixative (4% paraformaldehyde in 0.1 M phosphate buffer) for overnight fixation.

### *In vitro* electrophysiological recordings

Patch pipettes were pulled from borosilicate glass capillaries with filament (1.5 mm outer diameter and 1.1 mm inner diameter; Sutter Instruments) with a resistance of 2–3 MΩ. The pipette recording solution contained (in mM) 10 KCl, 130 K-gluconate, 1.8 NaCl, 0.2 EGTA, 10 HEPES, 2 Na-ATP, 0.2% Biocytin, pH 7.3 adjusted with KOH; 290-300 mOsm. Whole-cell recordings were made with Axopatch 700B amplifier (with Clampex 10.7 and Axoclamp1.1, Molecular Devices) using an upright microscope (Nikon Eclipse FN1, with ×40, 0.8 NA water immersion objective lens) equipped with differential interference contrast (DIC) optics. DIC images were captured with an Andor Zyla 5.5 sCMOS camera. All recordings were performed at 32 °C in ACSF bubbled with 95% O_2_ and 5% CO_2_. For AP parameter determination experiments, 1 μM CNQX (Sigma-Aldrich) was applied in the bath solution to eliminate EPSPs. Cells with <20 MΩ access resistance (continuously monitored) were accepted for analysis. Signals were low-pass filtered at 5 kHz and digitized at 20 kHz (Digidata 1550B, Molecular Devices). *In vitro* data analysis was performed with the help of Clampfit 10.7 (Molecular Devices) and Origin 8.6 (OriginLab Corporation).

### Immunohistochemistry, RNAscope *in situ* hybridization and confocal imaging

2weeks after IHKA injection animals were deeply anesthetized and transcardially perfused with ice-cold saline and then with 4% paraformaldehyde in 0.1 M PB (pH = 7.4). Brains and immersion fixed acute slices (patch clamp experiments) were cut into 60 μm thick sections in the coronal plane with a vibratome (Leica, VS1000s). On selected sections, immunoreactivities were tested: SATB1 (mouse, 1:500, SantaCruz) and GluR 2/3 (rabbit, 1:500, Millipore) were diluted in 0.1 M PB and incubated overnight at room temperature. For detection, fluorescent dye (Alexa488/Alexa594) conjugated donkey secondary antibodies (Jackson ImmunoResearch Labs) raised against the host species of primary antibodies were applied on the sections. To determine MC loss after SE, 10 sections (60 μm) per mouse (every third section) were collected starting from AP: -1.4 mm to AP: -3 mm from bregma and all SATB1 positive cells in the hilus were counted. In certain sections Immunohistochemistry was combined with RNAscope *in situ* hybridization as described previously[20]. Confocal images were taken with a Nikon Eclipse Ti2-E confocal microscope with 10x and 20x objectives.

### Object location and novel object memory test

Prior to the test, mice were habituated in the experimental room for 2 hours. During training phase, the mice were allowed to freely explore the environment for 10 minutes with the presence of two identical objects (LEGO blocks). During test phase (30 min after training), the mice were placed again in the presence of two objects. For OLM test the objects were the same, but one object was moved from the original location to a novel location. For NOR test one object was replaced with a novel object (LEGO blocks with different shape and color), but the location remained the original. Testing trials were video-recorded and analyzed using SolomonCoder software. Object exploration was quantified as the amount of time the mouse’s nose spent touching the object. The exploration times were expressed as a discrimination index (D.I. = (t_novel_ / t_total_) * 100.

## Results

### *Trpm4* is expressed in hilar MCs

To examine whether *Trpm4* is expressed in hilar MC cells, we used RNA scope *in situ* hybridization for its high sensitivity to detect even low level of expression. Unfortunately, immune-histochemical detection of TRPM4 protein is often controversial in the literature[21]. Using RNA scope in situ hybridization within the hippocampal formation, we found abundant *Trpm4* mRNA expression primarily in the hilar region (Figure 1 A-C). No expression was detectable in granule cells of the dentate gyrus. The high level of RNA scope signal in the hilus was consistent with the expression pattern found at the Allen Brain Atlas. To clarify which cell type in the hilus is expressing *Trpm4* we performed combined RNA scope and immunostaining using a novel molecular marker for MCs; Special AT-Rich Sequence Binding Protein 1 (SATB1). SATB1 is a transcriptional factor used as a neuronal marker in different brain areas[22] however its expression in MCs has never been shown. Double staining experiments using the MC marker GluR2/3[23] showed that 94.4±1.2 % of SATB1 positive cells were also GluR2/3 positive and 97.5±1.1 % of GluR2/3 positive cells were also SATB1 positive in the hilar region (Supplementary Figure 1). Since SATB1 stained exclusively MCs in the hippocampus contrary to GluR2/3 we therefore used this marker in our further experiments. Our combined *Trpm4* RNAscope and SATB1 immunostaining study clearly showed that the majority of SATB1 positive hilar MCs also express *Trpm4* (Figure1 A-G).

**Figure 1.**
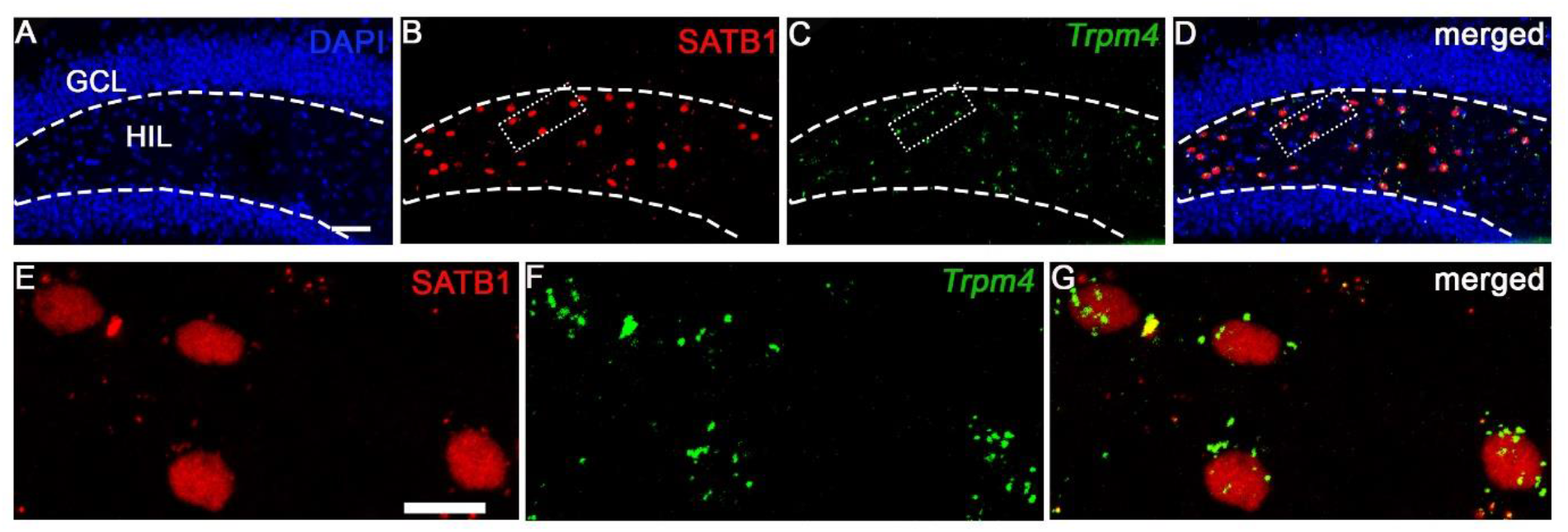
*Trpm4* is expressed in SATB1 positive hilar mossy cells. Representative 10x (A-D) and 60x (E-G) confocal images of immunofluorescence staining for STAB1 (red) and RNAscope labeling of *Trpm4* (green) from 3 independent experiments. Cell nuclei are stain with DAPI. Note, that *Trpm4* is specifically expressed in MCs but it is lacking from GCs. Scale bar 50 μm (upper row) and 5 μm (lower row).

**Supplementary Figure 1.**
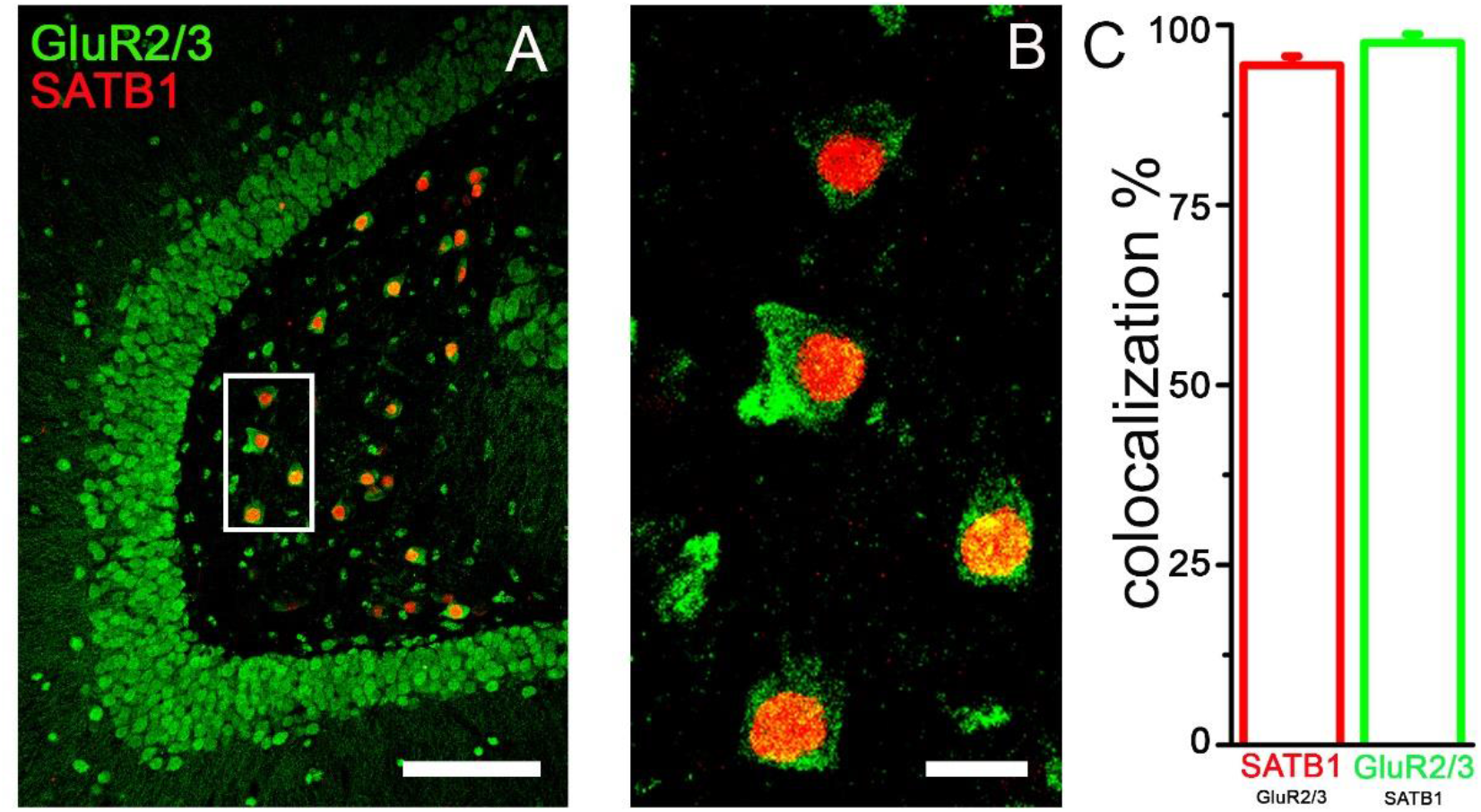
SATB1 and Glur2/3 are colocalized in the hilus. Representative 10x (A) and 60x (B) confocal images of double immunofluorescence staining for SATB1 (red) and GLuR2/3 (green). (C) Left, percentage of SATB1 positive neurons that express GLuR2/3. Right, percentage of GLuR2/3 positive neurons that express SATB1. n= 211 neurons from 2 mice. Scale bar 100 μm (left) and 5 μm (right).

### TRPM4 contributes to intrinsic electrophysiological properties of hilar MCs

In order to test whether TRPM4 is functionally active on hilar MCs we performed patch clamp recordings in acute brain slices containing the hippocampal formation. Randomly selected biocytin filled neurons were immunostained with the novel MC marker SATB1 (Supplementary Figure 2) for *post hoc* identification. Since there is no available selective and potent pharmacological blocker of TRPM4 - the most often used TRPM4 blocker, 9-phenantrol is only partially selective[24][25] – we used *Trpm4*^*-/-*^ mice to explore the role of the channel in MCs. Considering the timing of the increase in intracellular Ca^2+^ concentration relative to the neuronal AP[26] we assumed that a calcium activated cationic current might affect the repolarization phase of the AP. Indeed, when we compared AP parameters between WT and *Trpm4*^*-/-*^ MCs we found significantly greater after hyperpolarization (WT=-2.47±0.2 mV, *Trpm4*^*-/*-^=-4.34±0.64 mV, p=0.0049, two sample T-test) in neurons lacking TRPM4 indicating the absence of a depolarizing current during the late phase of the AP (Figure 2A). It has been shown previously that MCs are intrinsically active both *in vivo* and *in vitro*[4][23], therefore we explored whether this intrinsic activity is modulated by the action of TRPM4. Current clamp recordings revealed that the amplitude of the spontaneous EPSPs (WT=4.98±0.41 mV, *Trpm4*^*-/-*^ =3.92±0.23 mV, p=0.03, two sample T-test) and the frequency of the spontaneous APs (WT=1.33±0.22 Hz, *Trpm4*^*-/-*^=0.5±0.1 Hz, p=0,0019, two sample T-test) were both significantly lower in MCs lacking TRPM4 (Figure 2B-C). Finally, we challenged the cells with high concentration of glutamate (1 mM) mimicking excitotoxicity. Glutamate application increased the spontaneous AP frequency both in WT and *Trpm4*^*-/-*^ MCs however WT neurons reached depolarization block (AP amplitude decreased with more than 50%) more frequently than *Trpm4*^*-/-*^ cells (WT=6 out of 7 cells, *Trpm4*^*-/-*^ 1 out of 6 cells p=0.029, Fischer’s exact test).

**Figure 2.**
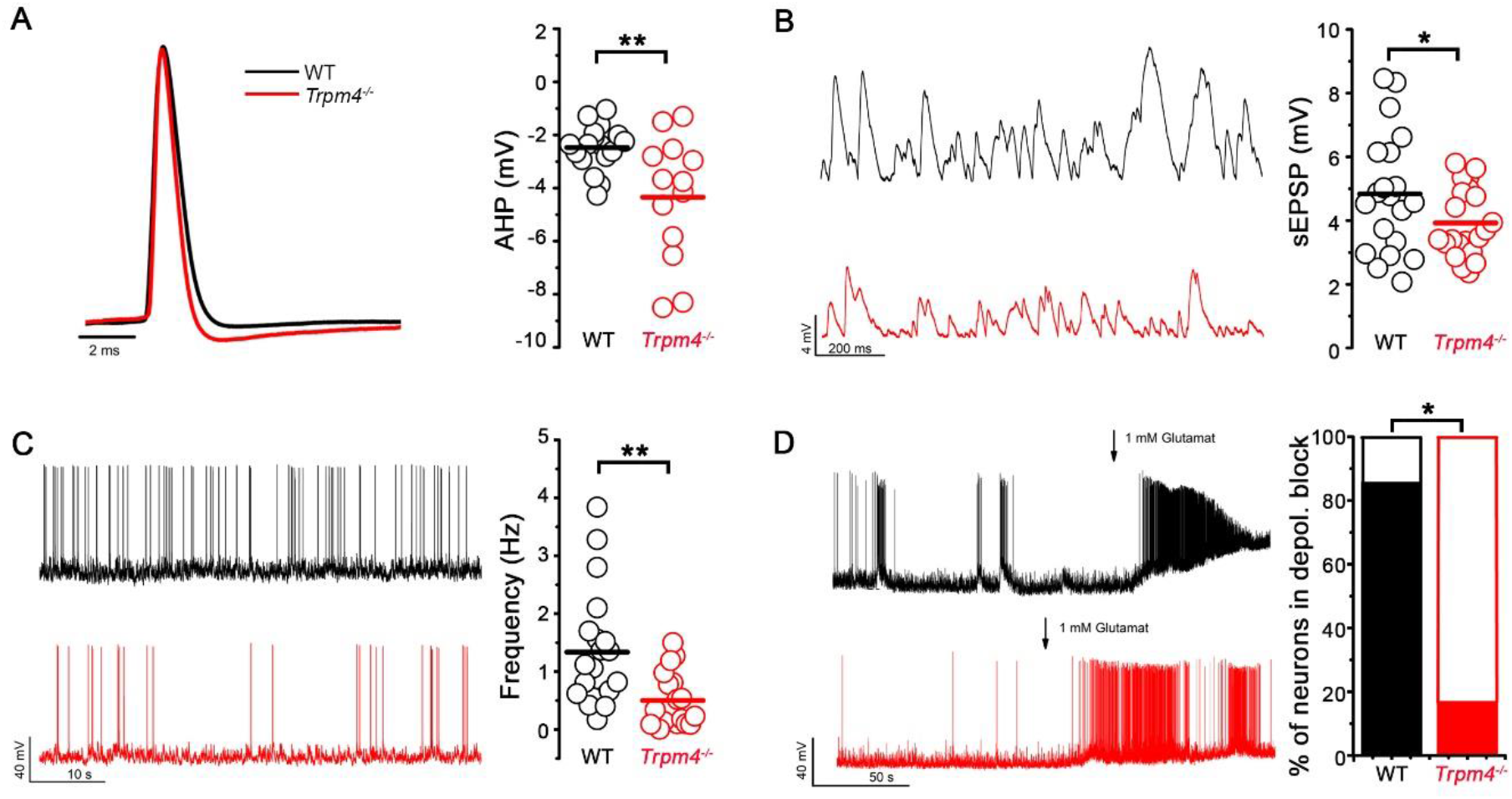
TRPM4 contributes to intrinsic electrophysiological properties of MCs. **(A)** Left, overlay of APs from WT (black) and *Trpm4*^*-/-*^ (red) MCs induced by current injection. Amplitudes of the representative APs are normalized to each other for better visibility. Right, statistics of AHPs from WT and *Trpm4*^*-/-*^ MCs. n=17 for WT and 13 for *Trpm4*^*-/-*^. **(B)** Left, representative traces of EPSPs from WT and *Trpm4*^*-/-*^ MCs. Right, statistics of AHPs from WT and *Trpm4*^*-/-*^ MCs. n=19 for WT and 21 for *Trpm4*^*-/-*^. **(C)** Left, representative spontaneous APs from WT and *Trpm4*^*-/-*^ MCs. Right, statistics of AP frequency from WT and *Trpm4*^*-/-*^ MCs. n=20 for WT and 19 for *Trpm4*^*-/-*^. **(D)** Left, representative current clamp recordings from WT and *Trpm4*^*-/-*^ MCs upon 1mM glutamate application. Right, statistics showing the percentage of neurons in depolarization block. n=7 for WT and 6 for *Trpm4*^*-/-*^. Data are presented as mean ± SEM. *p<0.05, **p<0.01.

**Supplementary Figure 2.**
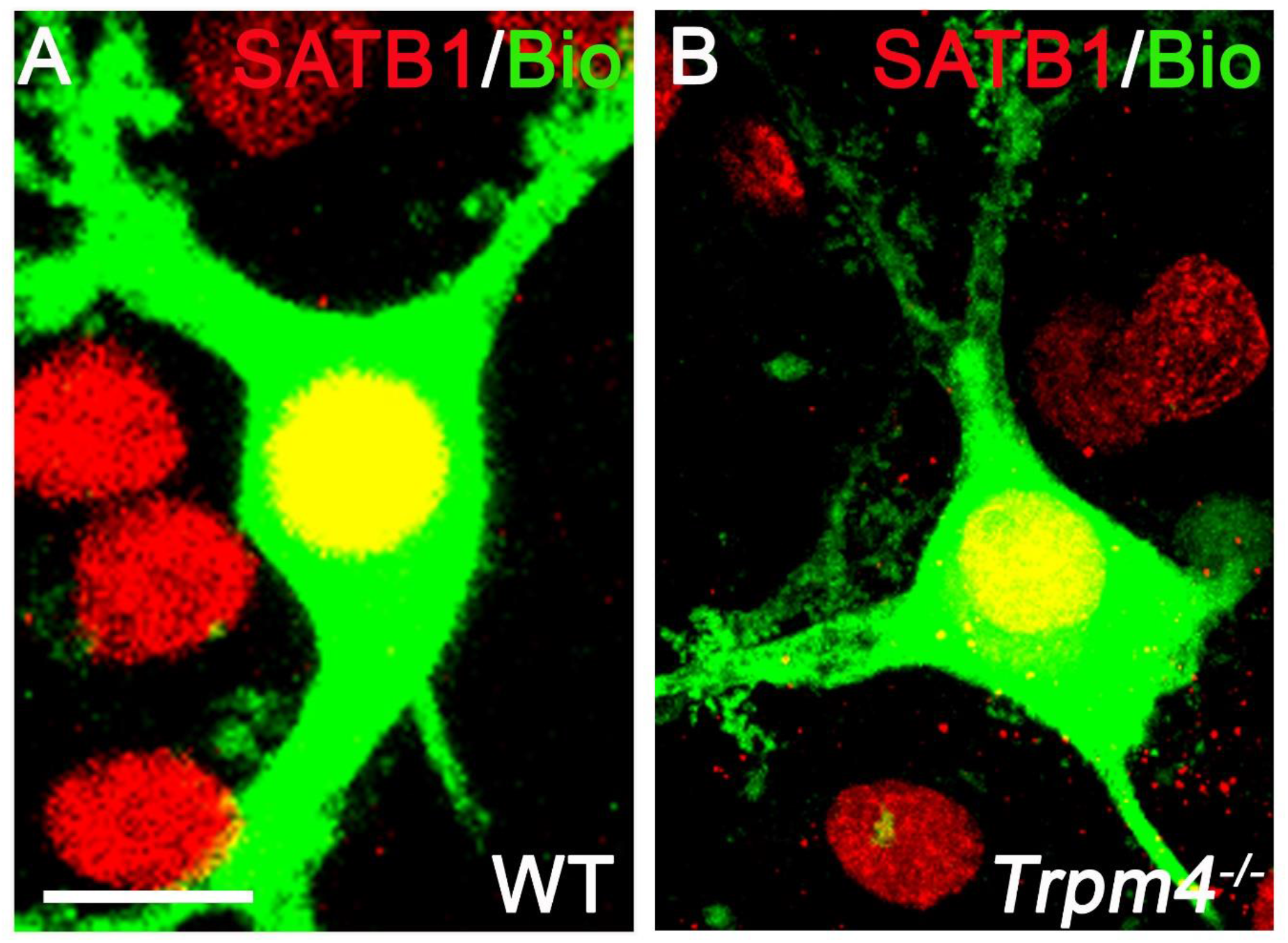
Patched neurons in the hilus are SATB1 positive. Representative confocal images of biocytin (green) filled WT **(A)** and *Trpm4*^*-/-*^ **(B)** MCs counterstained with SATB1. Note that the biocytin filled cells are also SATB1 positive (yellow). Scale bar 5 μm. Image of WT neuron (left) was modified from previous publication[6].

### *Trpm4*^*-/-*^ MCs are less vulnerable in experimental TLE compared to WT MCs

To examine MC vulnerability in experimental TLE we used the intrahippocampal kainic acid model (IHKA)[27]. KA was injected into the right dorsal hippocampus of WT and *Trpm4*^*-/-*^ mice. In our pilot experiments the concentration of KA (20 mM) often used in experimental model of TLE led to the practically complete elimination of MCs ipsilateral to the injection site (unpublished observation). Therefore, we decided to use a lower concentration (5 mM, 100 nL) to be able to evaluate the effect of SE specifically on MCs. To determine whether the lack of TRPM4 affected MC loss after SE, WT and *Trpm4*^*-/-*^ mice were sacrificed 14 days after KA injection. Brains were sectioned in the coronal plane and imunostained with SATB1 to evaluate MC loss in the hilus both ipsilateral and contralateral to the KA injection (Figure 3 A-C). From each mouse 10 sections were collected through the anterior-posterior length of the hippocampus and the average number of MCs per section was determined. These experiments revealed that SE resulted in significantly lower cellular survival rate in WT mice compared to *Trpm4*^*-/-*^ ipsilateral to KA injection (Figure 3 D) (WT_epileptic_= 18.93±5.46 %, *Trpm4*^*-/-*^_epileptic_= 53.72±3.84 %, p=0.00025, One-way ANOVA with Tukey’s post hoc test). Same tendency was present in the contralateral side although the difference was not significant (Figure 3 E).

**Figure 3.**
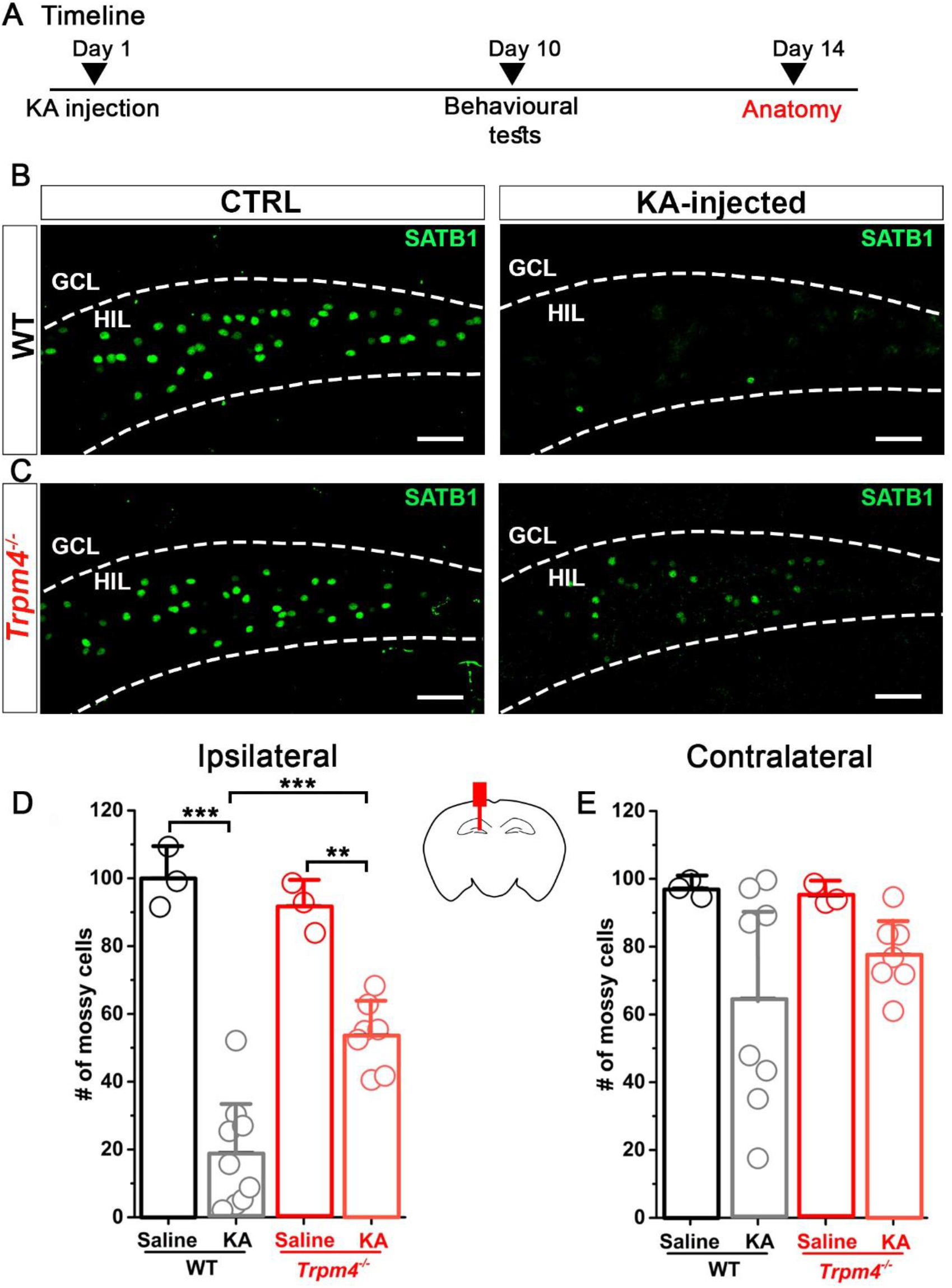
*Trpm4*^*-/-*^ MCs are more protected after KA injection compared to WT MCs. **(A)** Timeline of the experiments. Mice underwent KA or saline (ctrl) injection. 10 days later behavioral tests were performed to asses spatial and contextual memory (detailed in: Figure 5). At day 14 mice were sacrificed for histological studies. Representative 10x confocal images of SATB1 immunopositive neurons in the hilus from WT and **(B)** *Trpm4*^*-/-*^ **(C)** ctrl (left side) and KA (right side) injected mice. Statistics showing the number of MCs in WT and *Trpm4*^*-/-*^ mice ipsilateral **(D)** and contralateral **(E)** to the site of the injection. n=3 for WT_ctrl,_ 9 for WT_KA,_ 3 for *Trpm4*^*-/-*^_ctrl_ and 7 for *Trpm4*^*-/-*^_KA._ Cell numbers are normalized to the saline injected WT group. Scale bar 50 μm. Data are presented as mean ± SEM. *p<0.05, **p<0.01; ***p<0.001.

### *Trpm4*^*-/-*^ mice display decreased seizure susceptibility compared to WT mice

Next, we investigated whether the better survival of MC in *Trpm4*^*-/-*^ mice after SE also resulted in less frequent and severe spontaneous seizures during the chronic phase of the IHKA model. First we were interested whether we can detect interictal epileptic discharges (IED) preceding epileptic seizures in IHKA model[28]. WT and *Trpm4*^*-/-*^ mice underwent videoEEG recording 2 weeks after KA injection. Each mouse was monitored for 1 hour/day at the same time on 3 consecutive days (Figure 4 A-B). These experiments clearly showed a significantly increased number of IEDs in WT mice compared to *Trpm4*^*-/*^ mice (Figure 4 D) (WT=66.35±12.95 IED/hour, *Trpm4*^*-/-*^=24.45±9.1 IED/hour, p=0,017, Mann-Whitney test) indicating an increased hippocampal excitability. To examine the spontaneously occurring seizures we performed 24-hour videoEEG recordings 2-3 weeks after KA injection. The relatively low KA concentration we used in our experimental protocol resulted in only occasional seizures (1 seizure from 3 WT mice during 24 hours). Therefore, we decided to synchronize seizures with the injection of sub-threshold dose of KA intraperitoneally (i.p.). The sub-threshold dose of KA (5 mg/kg BW) resulted in no detectable seizures in healthy WT mice in line with previous observations[29] however, it induced both electrographic and behavioral seizures in IHKA mice (Figure 4 C). Both the number of seizures (WT=7.3±1.84 seizures/2hours, *Trpm4*^*-/-*^=1.5±0.74 seizures/2hours p=0,032, Mann-Whitney test) and the total seizure duration (WT=1186.1±409.83 sec/2hours, *Trpm4*^*-/-*^=336.5±231.13 sec/2hours p=0,088, Mann-Whitney test) were significantly lower in *Trpm4*^*-/-*^ mice in the first 2 hours following the i.p. KA injection (Figure 4 E-F). Interestingly, no difference was visible between the genotypes concerning the individual spikes (Figure 4 G). Furthermore, theta, alpha, beta and gamma power spectra were compared between epileptic WT and *Trpm4*^*-/-*^ mice but no significant changes were detected (Supplementary Figure 4).

**Figure 4.**
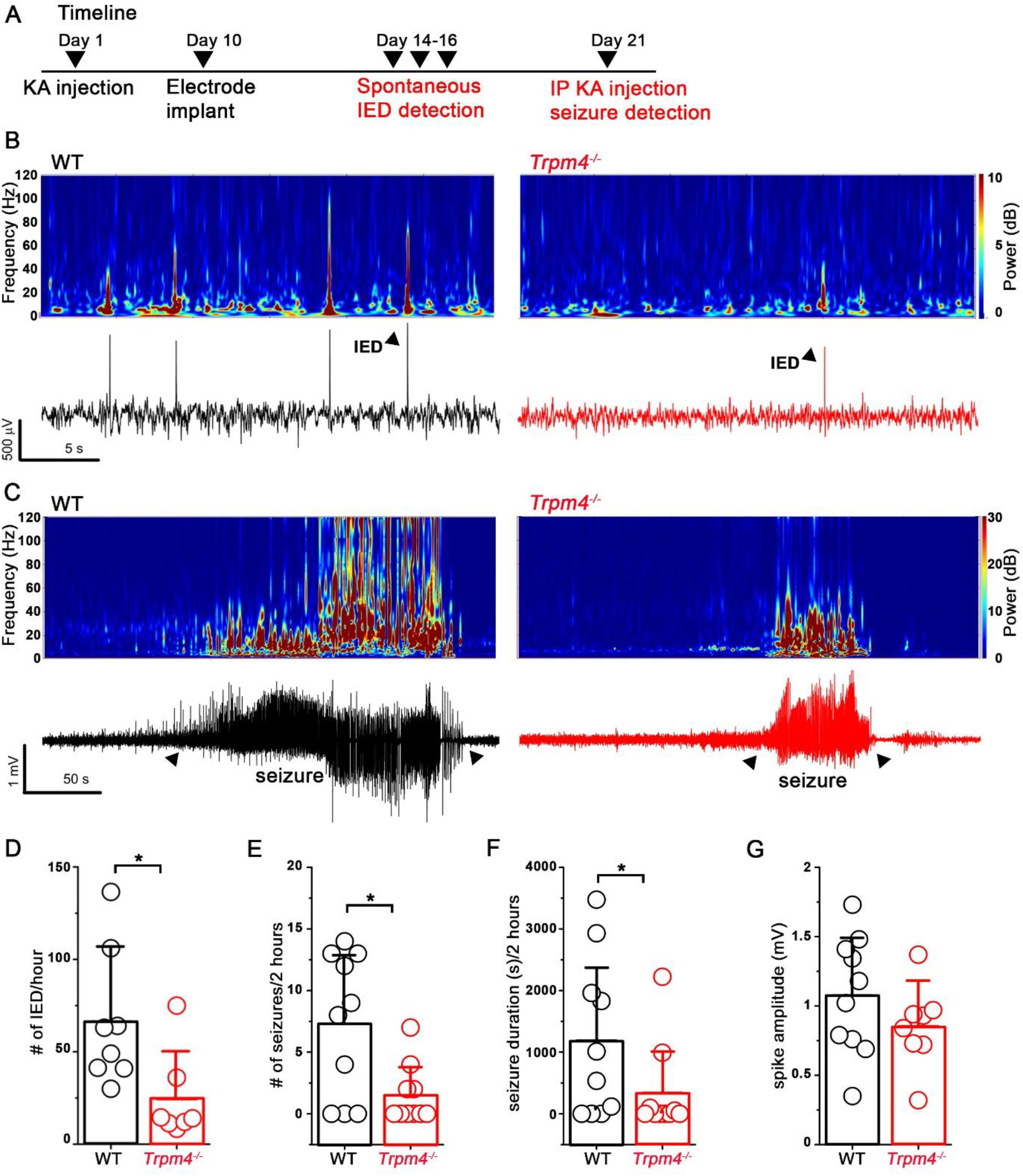
Increased seizure susceptibility in KA injected WT mice compared to *Trpm4*^*-/-*^. **(A)** Timeline of the experiments. Mice underwent KA or saline (ctrl) injection. 10 days later EEG electrodes were implanted. 14 days later mice underwent spontaneous IED detection, 21 days later seizures were induced with i.p. KA injection and detected with videoEEG. **(B)** Representative EEG recordings (lower panel) and corresponding spectrograms (upper panel) 14 days after KA injection in WT (black) and *Trpm4*^*-/-*^ (red) mice. Triangles indicates individual IED events. **(C)** Representative EEG recordings (lower panel) and corresponding spectrograms (upper panel) 21 days after KA injection in WT (black) and *Trpm4*^*-/-*^ (red) mice after KA injection. Triangles indicates seizures. **(D)** Statistics showing the number of spontaneous IED/hour 14 days after IHKA. n=8 for WT and 7 for *Trpm4*^*-/-*^. Statistics showing the **(E)** number of seizures, **(F)** seizure duration and **(G)** spike amplitude in 2 hours after i.p. KA injection. n=10 for WT and 10 for *Trpm4*^*-/-*^. Data are presented as mean ± SEM. *p<0.05.

**Supplementary Figure 4.**
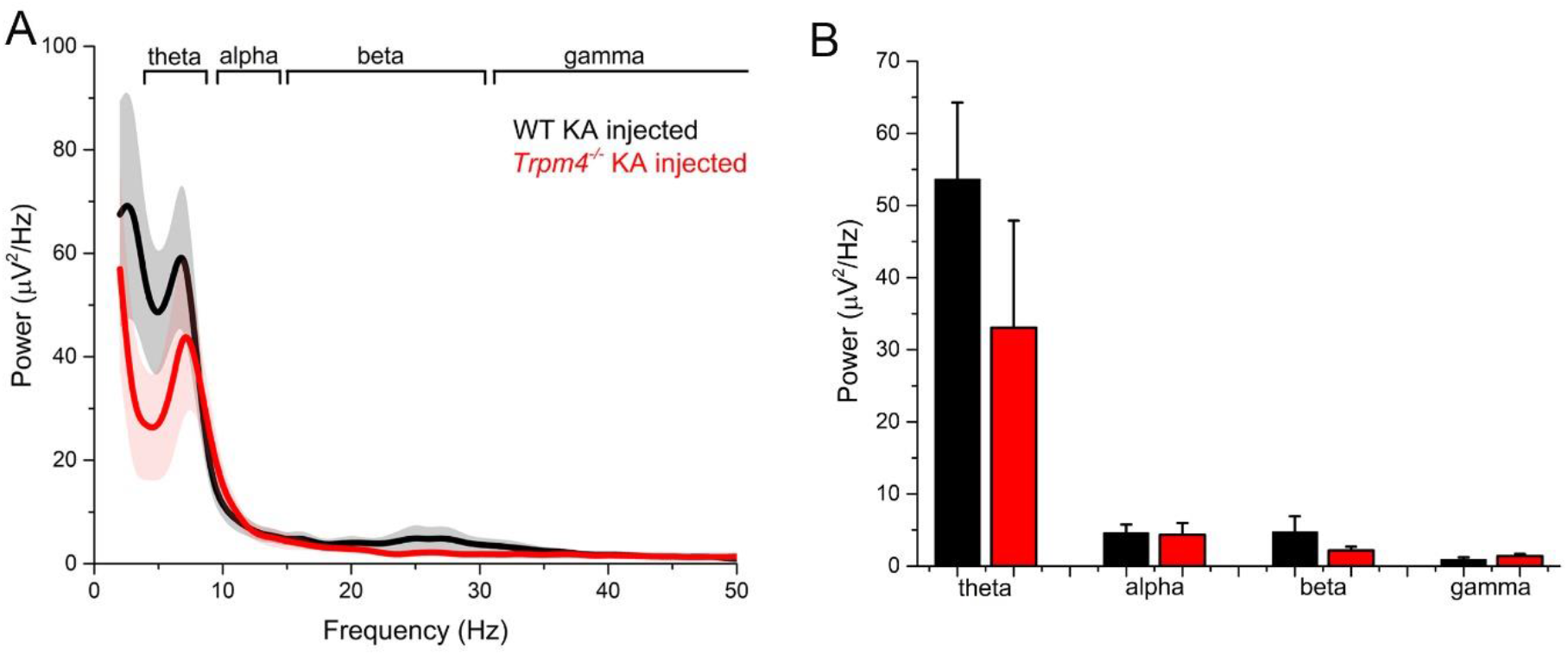
Power spectrum of epileptic WT and *Trpm4*^*-/-*^ mice is not different. **(A)** Power spectral density plot of epileptic WT (black) and *Trpm4*^*-/-*^ (red) mice during exploration. **(B)** Statistics showing theta (6 Hz), alpha (15 Hz), beta (25 Hz) and gamma (50 Hz) power in epileptic WT (black) and *Trpm4*^*-/-*^ (red) mice. n=6 for WT and 6 for *Trpm4*^*-/-*^ mice.

### TRPM4 deletion rescues memory impairment in chronically epileptic mice

To investigate the role of TRPM4 in memory impairment often seen in chronically epileptic mice[30] we performed object location memory (OLM) and novel object recognition (NOR) tests on control (saline injected) and epileptic mice on both genotypes (Figure 5A). Chronically epileptic WT mice had significant deficits in OLM test and were unable to distinguish the object in novel position from the unmoved object (Figure 5B-C). Interestingly, in epileptic *Trpm4*^*-/-*^ mice this significant deficit was not present as it is reflected in the discrimination index (WT_saline_=0.66±0.13 %, WT_epileptic_=0.45±0.05 %, *Trpm4*^*-/-*^_saline_=0.65±0.16 %, *Trpm4*^*-/-*^_epileptic_=0.54±0.13 %, p= WT_saline_ vs WT_epileptic_= 0.0071, *Trpm4*^*-/-*^_saline_ vs *Trpm4*^*-/-*^ _epileptic_=0.37, One-way ANOVA, Tukey’s post hoc test). On the contrary NOR test showed no deficit in chronically epileptic mice neither in WT nor in *Trpm4*^*-/-*^ mice (Figure 5E-F). Indicating that this type of contextual memory is not effected in our IHKA epilepsy model. Of note, these observations were not because of altered mobility between groups since object visits were not significantly different among them (Figure 5D,G).

**Figure 5.**
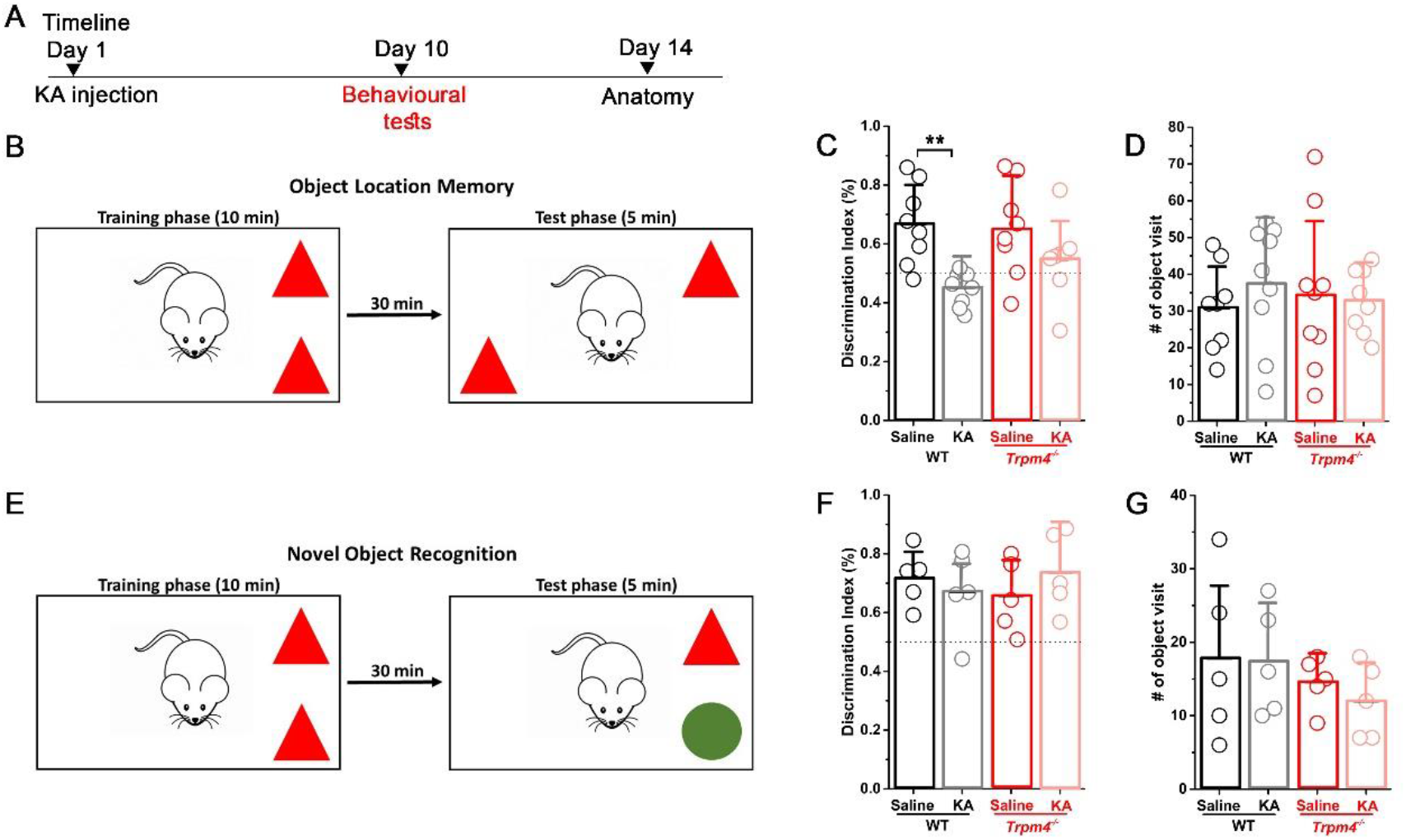
Epileptic WT but not *Trpm4*^*-/-*^ mice have impaired spatial memory. **(A)** Timeline of the experiments. Mice underwent KA or saline (ctrl) injection. 10 days later behavioral tests were performed to asses spatial and contextual memory. **(B)** OLM test schematic and timeline. Statistics showing discrimination index **(C)** and number of object visit **(D)** in WT and *Trpm4*^*-/-*^ mice. n=8 for WT_ctrl,_ 9 for WT_KA,_ 9 for *Trpm4*^*-/-*^_ctrl_ and 8 for *Trpm4*^*-/-*^_KA._ **(E)** OLM test schematic and timeline. Statistics showing discrimination index **(F)** and number of object visit **(G)** in WT and *Trpm4*^*-/-*^ mice. n=5 for WT_ctrl,_ 5 for WT_KA,_ 5 for *Trpm4*^*-/-*^_ctrl_ and 5 for *Trpm4*^*-/-*^_KA._ Data are presented as mean ± SEM. *p<0.05, **p<0.01; ***p<0.001.

## Discussion

In this study, we report that TRPM4 plays a pivotal role in MC loss during excitotoxic insults leading to TLE. At the cellular level, we showed for the first time that *Trpm4* is present in hilar MCs and as a Ca^2+^ -activated cationic current regulates the spontaneous activity of MCs as well as the repolarization of their AP in physiological conditions. In experimental TLE mice lacking TRPM4 showed reduced MC loss and seizure susceptibility during the chronic phase of epilepsy. Our finding that epileptic *Trpm4*^*-/-*^ mice show better performance in spatial memory tests further supports its role in MC vulnerability during seizures. In summary, our results provide evidence for the role of TRPM4 in MC excitability both in physiological and pathological conditions.

Expression pattern of TRPM4 in the brain is controversial because of the lack of selective antibodies. Several studies suggested the role of TRPM4 in neuronal function on different brain regions including the hippocampus[21][31][32] the prefrontal cortex[33] and the pre-Bötzinger[34] complex however, most of these publications used pharmacological tools with questionable selectivity[25] to investigate TRPM4 function. Recently, two studies proved the expression of *Trpm4* in brainstem pacemaker neurons using the ultrasensitive RNAscope *in situ* hybridization technique[35][36]. Our findings extend these results. To our best knowledge this is the first evidence that the Ca^2+^ activated cation channel *Trpm4* transcripts are present in hilar MCs. Furthermore, during these histological studies we introduced SATB1 as a novel molecular marker with exclusive selectivity towards MCs within the hippocampus.

What is the physiological role of TRPM4 in MCs? It has been shown previously that TRPM4-dependent membrane depolarization can contribute to the time course of the AP, as it was shown in cardiac myocytes, or it can support bursts of action potentials in neurons[13][14][17]. Furthermore, it can also mediate axonal and neuronal degeneration and glutamate mediated excitotoxicity in the animal model of experimental autoimmune encephalomyelitis[18]. In accordance, in our experiments we found an increased after-hyperpolarization in *Trpm4*^*-/-*^ MCs possibly indicating the lack of a Ca^2+^ activated depolarizing current at the late phase of the AP. Furthermore, MCs lacking TRPM4 displayed significantly decreased spontaneous AP firing frequency as well as decreased EPSP amplitude. Based on these observations we hypothesize that during excitatory postsynaptic events the increased calcium level activates TRPM4 and therefore amplifies EPSPs arriving onto MCs, and increases excitability of these cells. During an AP the depolarizing current via TRPM4 will counterbalance the end of the repolarization phase, which results in weak or no AHP commonly observed in MCs[3]. Of note, this would imply that TRPM4 is expressed both in the synapse and on the soma of MCs. Therefore it would be of great interest to conduct sub-cellular localization studies to precisely map TRPM4 expression. Moreover, one can speculate that the amplifier function of TRPM4 is an effective mechanism to enhance the inputs from the otherwise sparsely active GCs in physiological conditions, while during pro-epileptic insults it can worsen the excitotoxicity caused by the hyperexcited circuit. Indeed, challenging these cells with glutamate *Trpm4*^*-/-*^ MCs were less frequently entered into depolarization block compared to WT MCs.

A well-known experimental model for TLE is the direct injection of KA into the hippocampal formation (IHKA). This results in widespread cellular death among hilar MCs and the appearance of spontaneous seizures[27]. MCs are in a key position to regulate both the onset of epileptogenesis and the recurrent seizures during the chronic phase of epilepsy[6][5]. Our observation that MCs lacking TRPM4 are more protected during SE is probably the result of their reduced spontaneous activity, which is in line with previous observations that inhibiting MC during SE decreases MC and CA3 neuronal loss[9]. It is common view that the initial SE in TLE is followed by a seizure free period that precedes the development of spontaneous seizures and IED in between seizures[37]. In our experiments the reduced MC loss in *Trpm4*^*-/-*^ mice resulted in less frequent IEDs indicating that the overall excitability of the hippocampal circuit is reduced in these mice. Interestingly, despite the occurrence of IEDs in both genotypes after IHKA, spontaneously occurring seizures were rare. The reason for that most likely lies on the relatively low concentration of KA we used in our IHKA experiments. Nevertheless, this concentration range allowed us to more specifically investigate MC vulnerability between the two genotypes since the classically used higher dose practically killed more than 90 % of MCs in both genotypes, not surprisingly since MC are one of the most vulnerable cell types during epileptic insults[5]. Although spontaneous seizures were rare in our experimental protocol the seizure susceptibility clearly increased in IHKA mice. The subthreshold dose of i.p. KA injection did not resulte in seizures in naïve mice while it induced both electrographic and behavioral seizures in previously KA injected mice in both genotypes. Our findings that the induced seizures were less frequent and shorter in duration in case of *Trpm4*^*-/-*^ mice further support the protective role of TRPM4 elimination from MCs in epileptic conditions.

Finally, we found that the reduced loss of MCs in *Trpm4*^*-/-*^ mice during epilepsy results in better performance in spatial memory test. Interestingly, behavioral paradigms testing contextual novelty instead of spatial novelty has not changed in any of the genotypes during epilepsy further supporting the recent findings that MCs have an important role in spatial memory encoding[38][23].

Taken together we report here that *Trpm4* is expressed in hilar MCs and regulates their AP waveform and spontaneous activity in physiological conditions. Furthermore, in epilepsy TRPM4 contributes to MC loss and increase seizure susceptibility. Our results are especially important in the light of recent findings identifying TRPM4 as a promising drug target in cardiac arrhythmias[39] especially because it has been shown that this type of arrhythmia is often associated with seizures[40]. Thus TRPM4 might be a common regulator of Ca^2+^ dependent excitability both in cardiac myocytes and neurons. Better understanding of how TRPM4 regulates MC excitability may lead to novel strategies in seizure management and could highlight its clinical importance.

## Abbreviations

MC: mossy cells
GC: granule cell
IHKA: intrahippocampal kainic acid injection
AHP: after hyperpolarization
SE: status epilepticus
TLE: temporal lobe epilepsy
SATB1: Special AT-Rich Sequence Binding Protein 1
OLM: object location memory
NOR: novel object recognition
IED: interictal epileptic discharge

## Acknowledgements

The research was performed in collaboration with the Nano-Bio-Imaging and the Histology and Light Microscopy core facility at the Szentágothai Research Centre of the University of Pécs.

## Summary of statistics

Summary of statistics:
|  | n number | mean | sem | test | p value |
| --- | --- | --- | --- | --- | --- |
| **Figure 2 A** | WT=17<br>*Trpm4*<sup>-/-</sup>=13 | WT=-2.47<br>*Trpm4*<sup>-/-</sup>=-4.34 | WT=0.2<br>*Trpm4*<sup>-/-</sup>=0.64 | Two sample<br>T-test | 0.0049 |
| **Figure 2 B** | WT=19<br>*Trpm4*<sup>-/-</sup>=21 | WT=4.98<br>*Trpm4*<sup>-/-</sup>=3.92 | WT=0.41<br>*Trpm4*<sup>-/-</sup>=0.23 | Two sample<br>T-test | 0.03 |
| **Figure 2 C** | WT=20<br>*Trpm4*<sup>-/-</sup>=19 | WT=1.33<br>*Trpm4*<sup>-/-</sup>=0.5 | WT=0.22<br>*Trpm4*<sup>-/-</sup>=0.1 | Two sample<br>T-test | 0.0019 |
| **Figure 2 D** | WT=7<br>*Trpm4*<sup>-/-</sup>=6 | n.a. | n.a. | Fisher Exact<br>Test | 0.029 |
| **Figure 3 D** | WTsaline=3<br>WTKA=9<br>*Trpm4*<sup>-/-</sup>saline=3<br>*Trpm4*<sup>-/-</sup>KA=7 | WTsaline=100<br>WTKA=18.93<br>*Trpm4*<sup>-/-</sup>saline=91.8<br>*Trpm4*<sup>-/-</sup>KA=53.72 | WTsaline=5.15<br>WTKA=5.46<br>*Trpm4*<sup>-/-</sup>saline=4.2<br>*Trpm4*<sup>-/-</sup>KA=3.84 | One-way<br>Anova, Tukey's<br>post hoc | Wtsaline vs WTKA=<br>9.5123E-7 *Trpm4*<sup>-/-</sup><br>saline vs *Trpm4*<sup>-/-</sup><br>KA=0.0025 WTKA vs<br>*Trpm4*<sup>-/-</sup>KA=0.00025 |
| **Figure 3 E** | Wtsaline=3<br>WTKA=8<br>*Trpm4*<sup>-/-</sup>saline=3<br>*Trpm4*<sup>-/-</sup>KA=7 | Wtsaline=97.15<br>WTKA=64.62<br>*Trpm4*<sup>-/-</sup>saline=95.1<br>*Trpm4*<sup>-/-</sup>KA=77.68 | Wtsaline=1.43<br>WTKA=11.34<br>*Trpm4*<sup>-/-</sup>saline=1.7<br>*Trpm4*<sup>-/-</sup>KA=4.08 | One-way<br>Anova, Tukey's<br>post hoc | Wtsaline vs WTKA=<br>0.15<br>*Trpm4*<sup>-/-</sup>saline vs<br>*Trpm4*<sup>-/-</sup>KA=0.65<br>WTKA vs *Trpm4*<sup>-/-</sup><br>KA=0.65 |
| **Figure 4 D** | WT=8<br>*Trpm4*<sup>-/-</sup>=7 | WT=66.35<br>*Trpm4*<sup>-/-</sup>=24.45 | WT=12.95<br>*Trpm4*<sup>-/-</sup>=9.1 | Mann-Whitney<br>Test | 0.017 |
| **Figure 4 E** | WT=10<br>*Trpm4*<sup>-/-</sup>=10 | WT=7.3<br>*Trpm4*<sup>-/-</sup>=1.5 | WT=1.84<br>*Trpm4*<sup>-/-</sup>=0.74 | Mann-Whitney<br>Test | 0.032 |
| **Figure 4 F** | WT=10<br>*Trpm4*<sup>-/-</sup>=10 | WT=1186.1<br>*Trpm4*<sup>-/-</sup>=336.5 | WT=409.83<br>*Trpm4*<sup>-/-</sup>=231.13 | Mann-Whitney<br>Test | 0.044 |
| **Figure 4 G** | WT=10<br>*Trpm4*<sup>-/-</sup>=8 | WT=1.07<br>*Trpm4*<sup>-/-</sup>=0.85 | WT=0.13<br>*Trpm4*<sup>-/-</sup>=0.1 | Two sample<br>T-test | 0.22 |
| **Figure 5 C** | Wtsaline=8<br>WTKA=9<br>*Trpm4*<sup>-/-</sup>saline=9<br>*Trpm4*<sup>-/-</sup>KA=8 | Wtsaline=0.66<br>WTKA=0.45<br>*Trpm4*<sup>-/-</sup>saline=0.65<br>*Trpm4*<sup>-/-</sup>KA=0.54 | Wtsaline=0.13<br>WTKA=0.05<br>*Trpm4*<sup>-/-</sup>saline=0.16<br>*Trpm4*<sup>-/-</sup>KA=0.13 | One-way<br>Anova, Tukey's<br>post hoc | Wtsaline vs WTKA=<br>0.0071<br>*Trpm4*<sup>-/-</sup>saline vs<br>*Trpm4*<sup>-/-</sup>KA=0.37 |
| **Figure 5 D** | Wtsaline=8<br>WTKA=9<br>*Trpm4*<sup>-/-</sup>saline=9<br>*Trpm4*<sup>-/-</sup>KA=8 | Wtsaline=30.87<br>WTKA=37.44<br>*Trpm4*<sup>-/-</sup>saline=34.33<br>*Trpm4*<sup>-/-</sup>KA=32.87 | Wtsaline=11.87<br>WTKA=16.69<br>*Trpm4*<sup>-/-</sup>saline=20.8<br>*Trpm4*<sup>-/-</sup>KA=8.8 | One-way<br>Anova, Tukey's<br>post hoc | Wtsaline vs WTKA=<br>0.82 *Trpm4*<sup>-/-</sup>saline<br>vs *Trpm4*<sup>-/-</sup>KA=0.99 |
| **Figure 5 F** | Wtsaline=5<br>WTKA=5<br>*Trpm4*<sup>-/-</sup>saline=5<br>*Trpm4*<sup>-/-</sup>KA=5 | Wtsaline=0.719<br>WTKA=0.671<br>*Trpm4*<sup>-/-</sup>saline=0.657<br>*Trpm4*<sup>-/-</sup>KA=0.736 | Wtsaline=0.09<br>WTKA=0.14<br>*Trpm4*<sup>-/-</sup>saline=0.12<br>*Trpm4*<sup>-/-</sup>KA=0.13 | One-way<br>Anova, Tukey's<br>post hoc | Wtsaline vs WTKA=<br>0.93 *Trpm4*<sup>-/-</sup>saline<br>vs *Trpm4*<sup>-/-</sup>KA=0.75 |
| **Figure 5 G** | Wtsaline=5<br>WTKA=5<br>*Trpm4*<sup>-/-</sup>saline=5<br>*Trpm4*<sup>-/-</sup>KA=5 | Wtsaline=17.8<br>WTKA=17.4<br>*Trpm4*<sup>-/-</sup>saline=14.6<br>*Trpm4*<sup>-/-</sup>KA=12 | Wtsaline=11.27<br>WTKA=7.43<br>*Trpm4*<sup>-/-</sup>saline=3.5<br>*Trpm4*<sup>-/-</sup>KA=5.04 | One-way<br>Anova, Tukey's<br>post hoc | Wtsaline vs WTKA=<br>0.99 *Trpm4*<sup>-/-</sup>saline<br>vs *Trpm4*<sup>-/-</sup>KA=0.94 |

